# Cycle-by-cycle analysis of neural oscillations

**DOI:** 10.1101/302000

**Authors:** Scott Cole, Bradley Voytek

## Abstract

Neural oscillations are widely studied using methods based on the Fourier transform, which models data as sums of sinusoids. For decades these Fourier-based approaches have successfully uncovered links between oscillations and cognition or disease. However, because of the fundamental sinusoidal basis, these methods might not fully capture neural oscillatory dynamics, because neural data are both nonsinusoidal and non-stationary. Here, we present a new analysis framework, complementary to Fourier analysis, that quantifies cycle-by-cycle time-domain features. For each cycle, the amplitude, period, and waveform symmetry are measured, the latter of which is missed using conventional approaches. Additionally, oscillatory bursts are algorithmically identified, allowing us to investigate the variability of oscillatory features within and between bursts. This approach is validated on simulated noisy signals with oscillatory bursts and outperforms conventional metrics. Further, these methods are applied to real data—including hippocampal theta, motor cortical beta, and visual cortical alpha—and can differentiate behavioral conditions.

## Introduction

As a prominent feature of brain recordings, neural oscillations are frequently correlated to both pathologies (Uhlhaas and Singer 2010; Voytek and Knight 2015) and healthy behaviors such as movement, sleep, perception, and cognitive performance (Klimesch 1999; Massimini et al. 2004; Miller et al. 2007; Hanslmayr et al. 2007). Standard approaches for studying these oscillations are based on the Fourier transform, which decomposes a signal into component sinusoids. However, brain rhythms are not strictly sinusoidal nor stationary, as they come and go with varying amplitudes, frequencies, and waveforms (van Dijk et al. 2010; Jones 2016; Cole and Voytek 2017). Therefore, decomposition of the neural signal using the Fourier transform does not easily capture all of the interesting structure present in neural signals. This is suboptimal given that nonsinusoidal oscillatory features carry physiological information (Buzsáki et al. 1986; Hentschke et al. 2007; Pietersen et al. 2009; Mazaheri and Jensen 2010; Lewis et al. 2012; Belluscio et al. 2012; Lee and Jones 2013; Trimper et al. 2014; Sherman et al. 2016; Cole and Voytek 2017; Cole et al. 2017), and non-stationarities of low-frequency cortical oscillations may reflect different physiological processes (Peterson and Voytek 2017). Not properly accounting for these nonsinusoidal waveforms make conventional analyses susceptible to artifactual results, such as spurious phase-amplitude and cross-frequency coupling (Kramer et al. 2008; Lozano-Soldevilla et al. 2016; Gerber et al. 2016; Vaz et al. 2017; Cole et al. 2017; Scheffer-Teixeira and Tort 2016).

Methods used to analyze temporal properties of oscillations are also usually based on the Fourier transform. “Instantaneous” measures of oscillatory amplitude and frequency are widely used to estimate these time-varying properties of an oscillation of interest (Canolty et al. 2006; Samaha and Postle 2015). However, computation of such instantaneous features does not directly measure them in the raw (unbiased) recording, but always in a transformed version of the data, in which the signal is usually limited to a narrow sinusoidal frequency band (Bruns 2004). Such narrowband analyses can lead to misleading results, because they often overlook critical features present in the full power spectrum (Haller et al. 2018). This can lead to mischaracterizations of the data, such as an apparent increase in high frequency amplitude caused by a sharp transient, phase slips, and oscillatory frequency fluctuating within a single cycle (Kramer et al. 2008; Nelli et al. 2017).

Other complementary tools are therefore necessary to extract information from neural signals that Fourier-based analysis does not concisely capture. Matching pursuit is a tool for decomposing a signal using a dictionary of functions, and has been used to analyze transient components of brain signals (Ray et al. 2003; Chandran K S et al. 2016). However, this approach has only so far been applied with a basis of Gaussian-modulated sinusoids, and it is nontrivial to decide how to analyze the output to compare experimental conditions. Another approach, empirical mode decomposition (EMD), decomposes signals without forcing a basis function, such as the sinusoidal basis assumed in Fourier-based approaches (Pittman-Polletta et al. 2014). However, instabilities in the EMD algorithms can result in significant changes in results due to small changes in hyperparameters. Furthermore, interpreting the resultant EMD components can be difficult.

Few analytic approaches have been designed specifically for characterizing the time-domain waveform shape of brain oscillations. Recently, however, two algorithms were developed to extract the waveform of the *prominent* oscillator in a neural signal (Jas et al. 2017; Gips et al. 2017), but these approaches do not capture changes in waveform shape within a recording. We recently reviewed that waveform shape is diverse across the brain and relates to physiology, pathology, and behavior (Cole and Voytek 2017). The hippocampal theta rhythm (Buzsáki et al. 1985), cortical slow oscillation (Amzica and Steriade 1998), and the mu rhythm (Pfurtscheller et al. 1997) are particularly known to have stereotyped nonsinusoidal waveforms. There are a wide variety of circuit activation patterns for oscillators of each frequency (Womelsdorf et al. 2014), and the specifics of these dynamics may relate to the temporal dynamics of a single cycle of the recorded oscillation, or its waveform shape. Therefore, differences in waveform shape may hint at differences in the parameters, conditions, or even qualitative mechanisms of the oscillatory generator. One potential, or at least intuitive, interpretation of waveform shape is that sharper oscillatory extrema may be produced by more synchronous neural activity (Sherman et al. 2016; Cole et al. 2017). This may be caused by excitatory synaptic currents occurring relatively simultaneously in a cortical region and integrating in the local field to yield a sharp waveform, whereas those same currents, more spread out in time, will result in a smoother local field potential.

Here we present a time-domain approach, complementary to traditional frequency-domain analysis, designed to characterize nonsinusoidal and transient brain rhythms in order to help quantify information missed in conventional, Fourier-based neural signal processing. This novel framework analyzes oscillatory features on a cycle-by-cycle basis, and the necessary code is made open-source as a user-friendly Python toolbox^1^ with extensive tutorials and examples^2^. For each cycle, amplitude and period (frequency) are quantified, as are its waveform symmetries. In contrast to the instantaneous features cited above, cycle-by-cycle measures are computed from straightforward features rather than relying on transforms that assume a sinusoidal structure. Importantly, the output also specifies whether the oscillation of interest is present or absent in the signal during each “cycle” period, as it is unlikely that the oscillator is present throughout the whole duration of the signal (Fransen et al. 2015; Jones 2016). This is important, as estimates of oscillatory features are meaningless if no oscillation is evident.

## Methods

All methods described here are available in the open-source package ‘neurodsp’, available at https://github.com/voytekresearch/neurodsp. and all Python code to replicate the figures in this paper are shared at https://github.com/voytekresearch/Cole_2018_cyclebycycle. All tests are nonparametric, such that correlations are Spearman, two-sample unpaired tests are Mann-Whitney U. and two-sample paired tests are Wilcoxon Signed Rank.

### Segmentation of signal into cycles

The first step in characterizing individual oscillatory cycles in a neural signal is to split the entire recording into cycles. We first identify putative peaks and troughs throughout the recording. The raw data (Figure 1A) is optionally filtered with a broad bandpass filter to remove slow transients or high frequency activity that may interfere with peak and trough identification (Figure 1B). Next, the signal is bandpass filtered in the frequency band of interest, ideally verified from the power spectral density (Haller et al. 2018), and the time points of the rising and decay zero-crossings are identified (Figure 1C). The minima and maxima between these zero-crossings are declared as putative peaks and troughs (Figure 1D). Finally, the midpoints of the rise and decay flanks are computed by finding the time point at which the voltage is halfway between the peak and trough voltage (Figure 1E). Together, the extrema and flank midpoints can be used to estimate a phase time series (Figure 1F, black) by interpolating between their theoretical phases (e.g., peak has phase 0, the decay midpoint has phase π/2, etc.). This estimate deviates from the instantaneous phase commonly computed by bandpass filtering the signal in the oscillatory frequency band and applying the Hilbert transform (Figure 1F, red) but it more closely matches intuitions for the locations of an oscillatory peak and trough, which can be skewed when a nonsinusoidal oscillator is filtered in a narrow frequency band.

**Figure 1.**
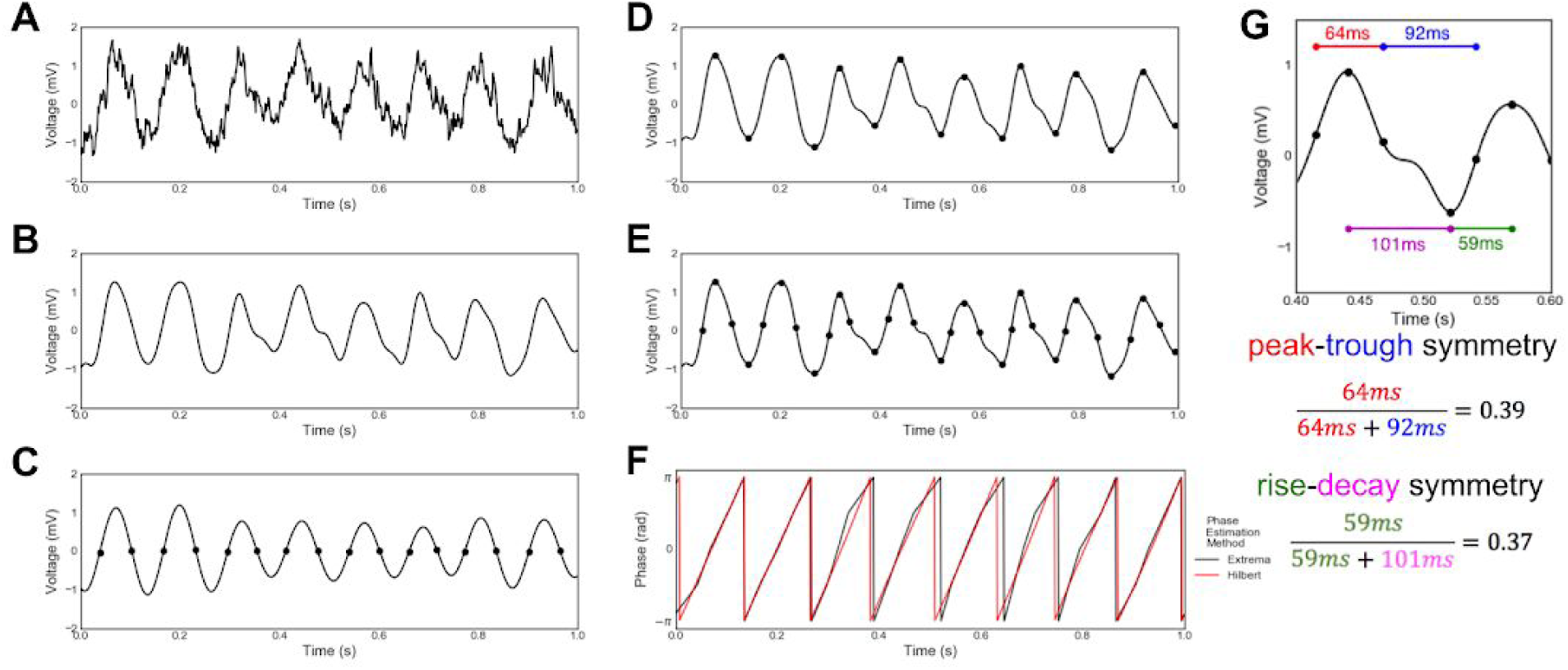
Decomposing a neural signal into individual cycles. **(A)** Example raw field potential recording from the CA1 layer of rat hippocampus. **(B)** The raw signal is broadly bandpass filtered (1-25 Hz) in order to remove high frequencies while preserving underlying theta waveform shape. **(C)** Zero-crossings are found after the signal has been bandpass filtered in the theta frequency range (4-10 Hz). **(D)** Peaks and troughs are found in the broad-bandpass filtered signal by finding the relative maxima and minima between the zero-crossings found in the theta-filtered signal **(C)**. **(E)** Flank midpoints are determined by locating the time points at which the voltage is halfway between the adjacent peak and trough voltages. These points denote the boundaries between peak and trough. **(F)** Instantaneous phase is estimated by interpolating between extrema and flank midpoints (black). This waveform-based phase estimate is contrasted against the analytic phase estimated using the more standard, Fourier-based filtering and Hilbert transform approach (red). Discrepancies between the two approaches are notable. **(G)** Demonstration of computed symmetry of a single cycle. The rise-decay symmetry is computed as the fraction of the period that the cycle is in the rise phase. Similarly, the peak-trough symmetry is computed as the fraction of the period between the rise midpoint and the subsequent decay midpoint (i.e., the peak).

### Cycle feature computation

After the signal is segmented into cycles. each cycle is characterized with a few basic measures. In all analvses. cycles are chosen to start and end at consecutive peaks. The amplitude of the cycle is computed as the average voltage difference between the trough and the two adjacent peaks. The period is defined as the time between the two peaks. Rise-decay symmetry (rdsym) is the fraction of the period that was composed of the rise time (Figure 1G; rdsym = 0.5 is a cycle with equal durations of rise and decay). Peak-trough symmetry (ptsym) is the fraction of the period, encompassing the previous peak and current trough, that was composed of the peak (Figure 1G; ptsym = 0.5 is a cycle with equal durations of peak and trough). The distributions of these features can be computed across all cycles in a signal in order to compare oscillation properties in different neural signals (Figure 2).

**Figure 2.**
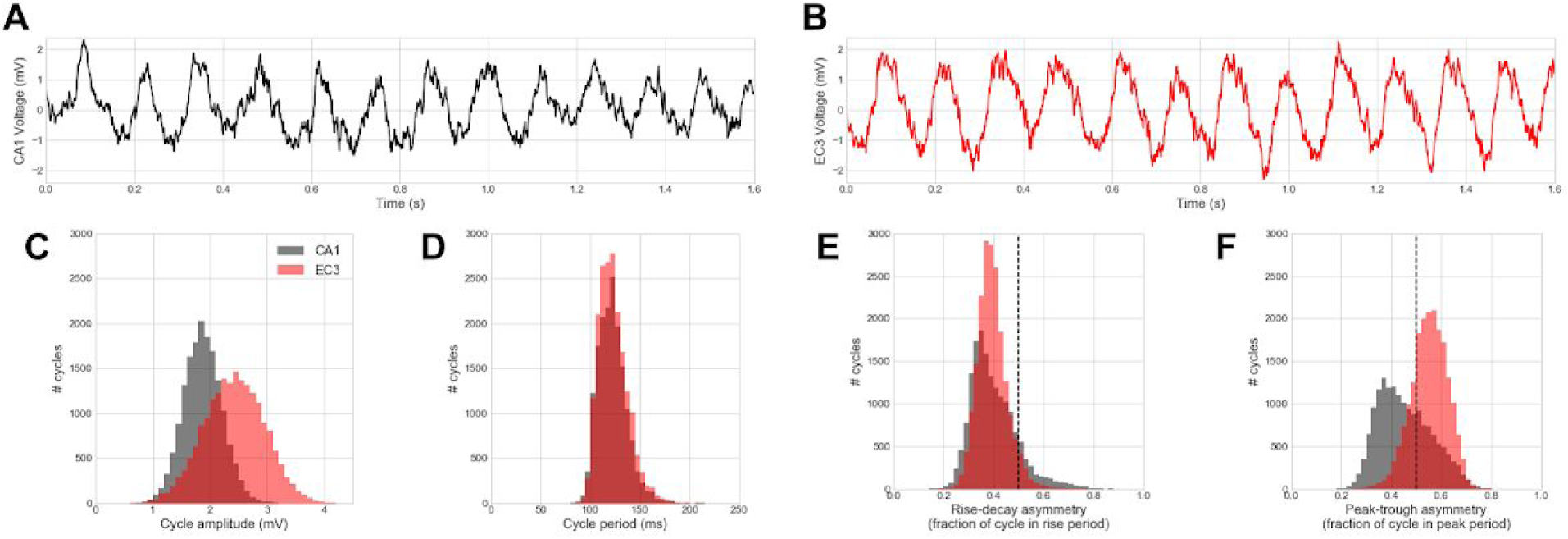
Distributions of cycle features differ between theta oscillations recorded in the CA1 pyramidal layer and layer 3 of the entorhinal cortex (EC3) **(A,B)** Raw voltage recording of theta oscillations in **(A)** CA1 and **(B)** EC3 **(C-F)** Distributions of cycle: **(C)** amplitude, **(D)** period, **(E)** rise-decay symmetry, and **(F)** peak-trough symmetry for CA1 (black) and EC3 (red) theta oscillations. Vertical dashed lines at 0.5 in E,F note relative rise-decay and peak-trough symmetry, respectively. Note that EC3 oscillations are **(C)** relatively higher amplitude and **(F)** have relatively longer peaks than CA1 oscillations, while the two show little difference in **(D)** their period (frequency) or **(E)** rise-decay asymmetry. This emphasizes the uniqueness of information provided by these different measures and how they might be used to uncover new physiological insight.

### Determination of oscillatory burst periods

After computing features for each cycle, an algorithm is applied to determine whether each cycle is part of an oscillatory burst (Figure 3). Oscillatory bursts are identified as time periods in which consecutive cycles in the time series had similar amplitudes, similar periods, and rise and decay flanks that are predominantly monotonic. To test this, three additional features are computed for each cycle. First, the amplitude consistency of a cycle is quantified as the relative difference in the rise and decay voltage (0.5 corresponds to one flank being 2 times longer than another, 1.0 corresponds to the flanks being equal length, etc.). The minimum value is taken after computing this measure for each pair of adjacent flanks that includes the cycle’s rise or decay flanks. The period consistency feature is computed as the maximal relative difference between the cycle’s period and the period of the adjacent cycles (0.5 corresponds to the previous or subsequent period being twice or half the length of the current cycle). The third and final feature, monotonicity, is the fraction of instantaneous voltage changes (difference between consecutive samples) that are positive during the rise phase and negative during the decay phase (0.8 corresponds to 20% of the voltage time series going in the opposite direction of the current flank).

**Figure 3.**
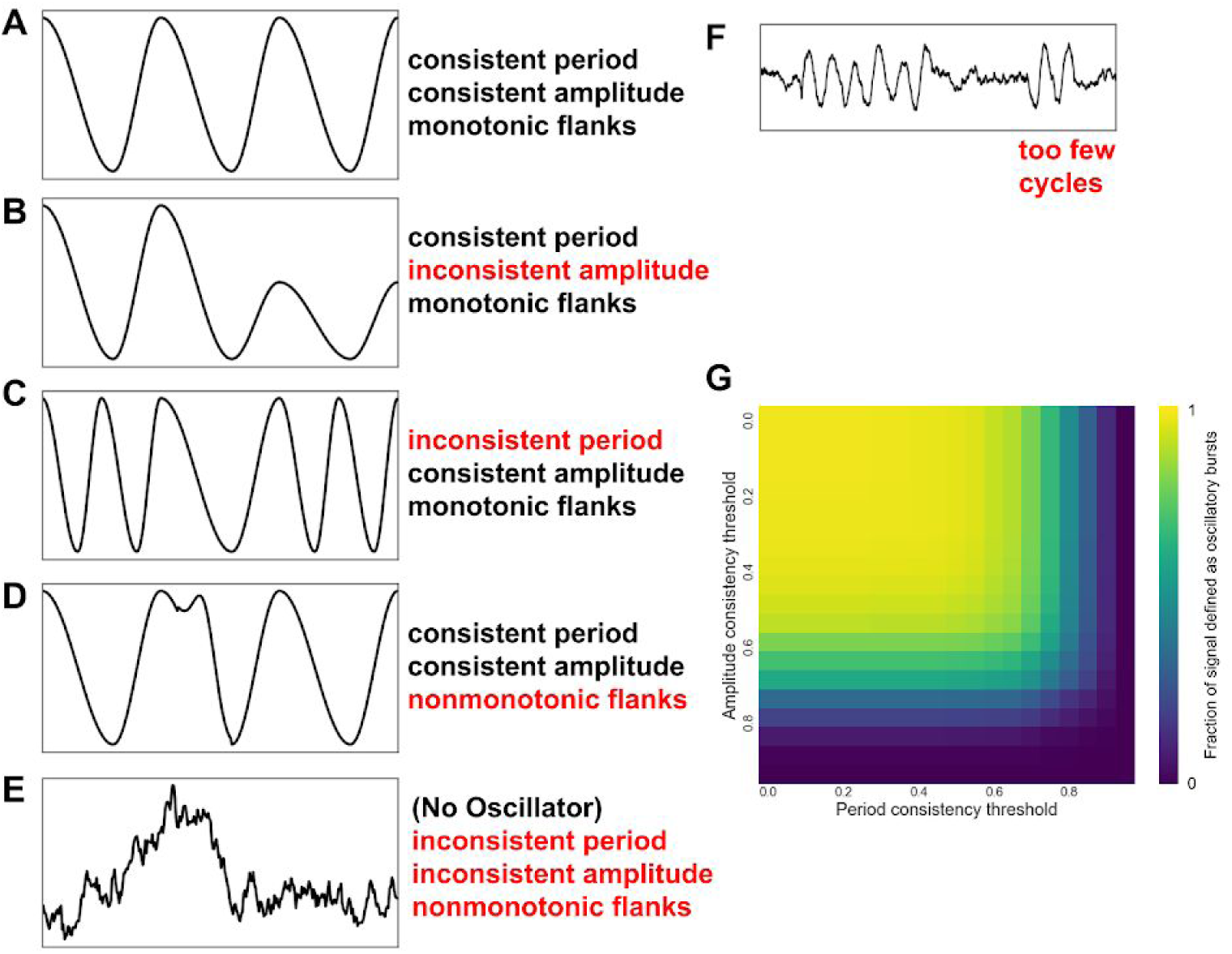
Determining oscillatory bursts in a neural signal. **(A-F)** In order for cycle-by-cycle measures to be meaningful, first a cycle must be determined to be a part of an oscillation. To determine this, its amplitude and period must be about the same as the adjacent cycles. Additionally, the rise and decay flanks should be mostly monotonic. According to these criteria: **(A)** passes all these requirements; **(B)** fails because the amplitudes are inconsistent; **(C)** fails because the period is inconsistent; **(D)** fails because the decay flank is considerably nonmonotonic, and; **(E)** fails because the peaks and troughs found in this noisy signal would violate all of the aforementioned requirements. The second burst in **(F)** would fail to be considered oscillatory because there are too few cycles present (minimum is 3 cycles). **(G)** Because these decision points are flexible, we examined how they interact. The fraction of cycles marked as oscillating decreases as the amplitude and period consistency requirements are set to be more strict (closer to 1).

Thresholds are set for each of these three features, as well as an optional threshold for the relative oscillation amplitude, in order to match the user’s judgment for what data segments should be defined as oscillations. If the consistency and monotonicity features are all above their respective thresholds, then they are marked as oscillating periods. The assignments are updated to require at least three consecutive cycles to be marked as an oscillating period (Figure 3F). In the current study, cycles were only analyzed that were defined as oscillating periods. For all hippocampal and simulated recordings, the thresholds used were those that maximized the Fβ score as shown in Figure 4H (amplitude consistency threshold = 0.6, period consistency threshold = 0.75, monotonicity threshold = 0.8). The Fβ score is used to trade off between precision and recall, where β is the relative importance weight between precision and recall (see below). For the motor cortical beta and visual cortical alpha analyses, thresholds were tuned in order to improve oscillatory burst detection because the signal-to-noise ratio (SNR) of the oscillator is relatively lower in these signals (amplitude consistency threshold = 0.3, period consistency threshold = 0.5, monotonicity threshold = 0.6). Both analyses also required cycles to be above the twentieth percentile of amplitude relative to the whole recording in order to be considered oscillatory bursts.

**Figure 4.**
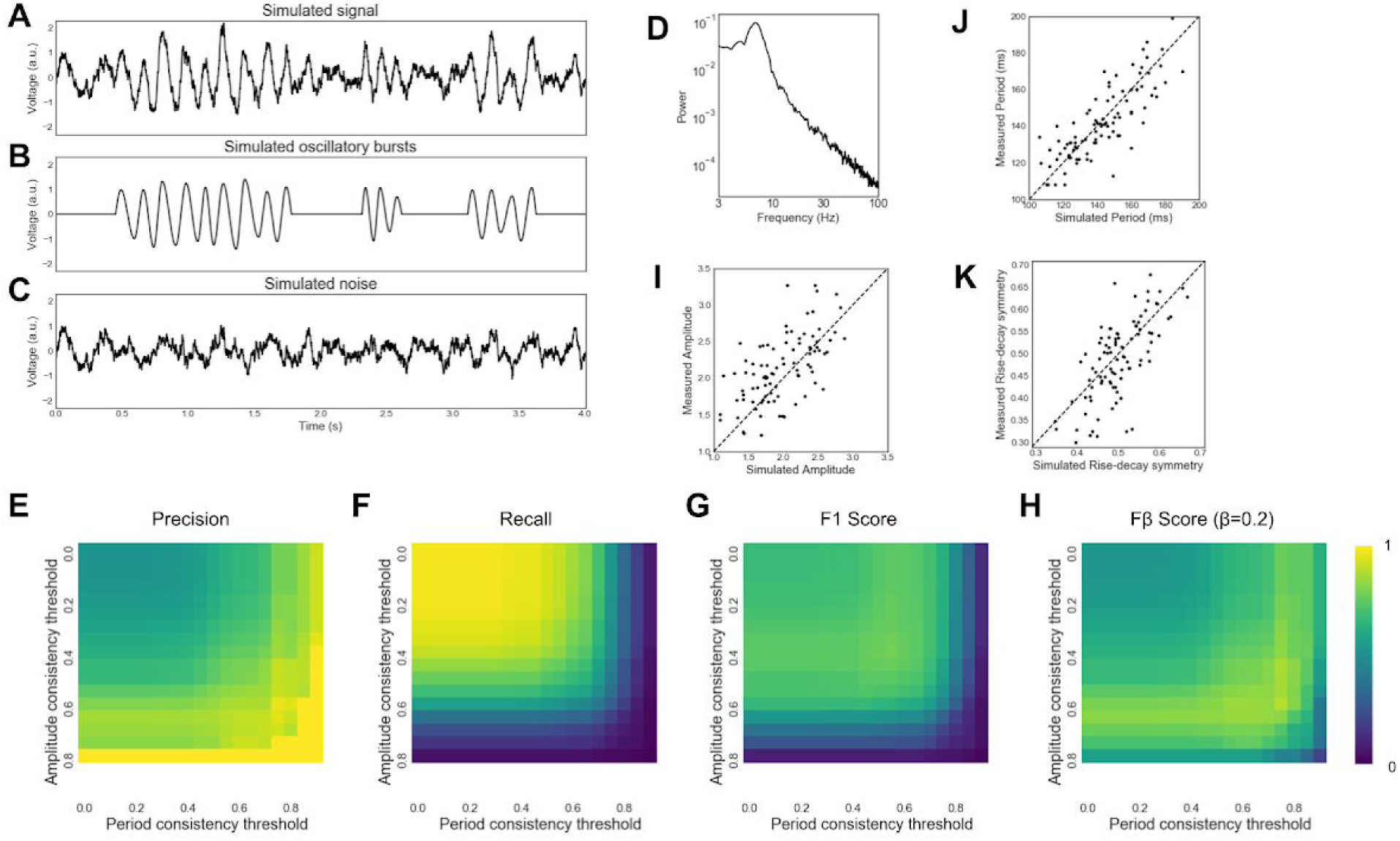
Validating cycle-by-cycle analysis with simulated data. **(A-C) (A)** A neural signal was simulated by adding, **(B)** a non-stationary bursting oscillation to, **(C)** brown noise. **(D)** Power spectral density of the simulated neural signal contains a peak at 7 Hz and a 1/f^2^ aperiodic process. **(E-H)** Accuracy of the algorithm used to determine if a cycle is in an oscillation. Amplitude and period consistency requirements were varied and accuracy was measured by (E) precision, **(F)** recall, **(G)** accuracy (F1 score), and **(H)** effectiveness (Fβ score; β = 0.2). **(I-K)** There was a positive correlation between a cycle’s simulated **(I)** amplitude, **(J)** period, and **(K)** rise-decay symmetry and their measured values. These results show that our approach, using the decision parameters outlined, are successfully capturing the ground-truth data.

### Simulation of oscillatory bursts with noise

Voltage time series were simulated to have properties of real neural recordings (Figure 4A). Specifically, a bursting oscillatory process (Figure 4B) was generated in a specified frequency range and added to brown (1/f^2^) noise that was highpass filtered with a cutoff frequency of 2 Hz (Figure 4C). The relative power of the oscillation and noise were controlled by scaling the amplitude of the brown noise using an SNR parameter. The bursting oscillator was defined by probabilities for entering and leaving an oscillatory regime. Furthermore, the amplitude, period, and rise-decay symmetries were randomly generated for each cycle by sampling from a Gaussian distribution defined by adjustable means and standard deviations. To account for the positive correlation of features between cycles within a burst (Figure 7), an additional standard deviation term was added so that a different mean feature value was computed for each burst.

In order to compare cycle-by-cycle measures with instantaneous amplitude and frequency (Figure 5), trials were simulated in which a 10 Hz oscillation was induced after an event. Brown noise was simulated throughout the trial (-1 to 2 seconds), and a bursting oscillator was simulated from 0 to 2 seconds in the same manner as above, except that the amplitude, period, and symmetry of oscillations were kept constant. Three conditions of 100 trials each were simulated for assessing amplitude measures. The “higher amp” condition induced alpha oscillations with a 20% greater amplitude than in the “baseline” condition. The “longer bursts” condition was half as likely to end a burst and 50% more likely to enter a burst compared to the other conditions. Similar trials were simulated for comparing cycle-by-cycle and instantaneous measures of frequency. Four conditions were simulated: “baseline”, “longer bursts”, “faster”, and “nonsinusoidal”. In the “faster” condition, an 11 Hz oscillation was simulated instead of a 10 Hz oscillation. In the “nonsinusoidal” condition, the oscillations simulated had a constant rise-decay symmetry value of 0.2 (other conditions: 0.5).

**Figure 5.**
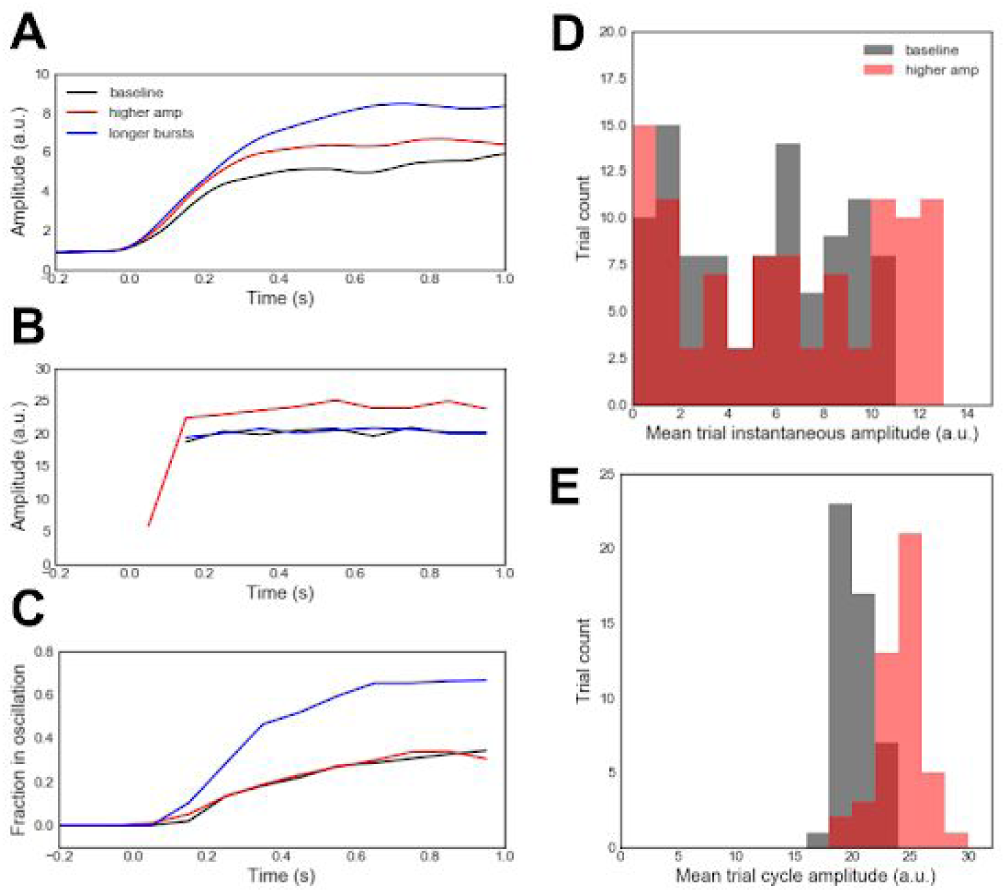
Comparison of cycle-by-cycle and instantaneous amplitude analysis on simulated event-related data. Three different event-related oscillatory changes were simulated for an event occuring at 0.0 seconds: (1) a “baseline” condition in which short, 10 Hz oscillatory bursts were injected into the data after a simulated event onset; (2) a “higher amplitude” condition in which oscillations had 20% higher amplitude than in the “baseline” condition, and; (3) a “longer bursts” condition wherein bursts were half as likely to end and 50% more likely to begin compared to the other conditions. **(A)** Event-related instantaneous amplitude changes, using conventional Fourier and Hilbert approaches, were averaged across three types of trials: “baseline” (black), “higher amp” (blue), and “longer bursts” (red). **(B)** Same as **(A)**, but using cycle-by-cycle approaches introduced here. Amplitude estimates were averaged across trials and binned at 100 ms intervals. This amplitude estimate is not defined when no oscillating bursts are detected, hence the lack of an average before the event. **(C)** Same as **(B)**, but showing the fraction of cycles that were defined as oscillating for each condition, binned at 100 ms intervals. Note that this approach is not standard for Fourier-based analyses. **(D,E)** Distributions of the average **(D)** instantaneous amplitude and **(E)** cycle amplitude between 500 ms and 1000 ms for each trial in the “baseline” (black) and “higher amp” (blue) conditions. Note that conventional Fourier and Hilbert approaches only weakly identify event-related amplitude increases as such **(A,D)** while misclassifying longer bursts as increased amplitude **(A)**. This is the result of averaging across trials. This mischaracterization is mitigated by the cycle-by-cycle approach introduced here, which successfully differentiates event-related changes in amplitude **(B,E)** from changes in burst duration **(C)**.

### Instantaneous amplitude and frequency computation

Instantaneous measures of amplitude and frequency were computed using common methods and applied to the simulated alpha oscillations described in the previous paragraph. Signals were first bandpass filtered (8-12 Hz) and then the Hilbert transform was applied. The magnitude of the resultant time series was computed to obtain the instantaneous amplitude estimate, and the angle was computed to obtain the instantaneous phase estimate. Instantaneous frequency was computed from the instantaneous derivative of the phase time series, and then median filtered at multiple time scales as previously described (Samaha and Postle 2015).

### Hippocampal theta recordings

Local field potential recordings from the CA1 pyramidal cell layer of the hippocampus and layer 3 of the entorhinal cortex (EC3) were obtained from the “hc3” dataset on the Collaborative Research in Computational Neuroscience (CRCNS) database (Mizuseki et al. 2014; Teeters et al. 2008). A recording was chosen in which CA1 and EC3 were recorded simultaneously, and for which position data was also available (rat ec014, session 440, the deepest contact on shanks 3 and 11) in which the rat was performing an alternation task in a T-maze with wheel running (Pastalkova et al. 2008). Another recording of CA1 theta oscillations from the same database was used in Figure 7. For more information on the data collection, see Mizuseki et al. (2014). The speed of the rat was estimated for each cycle by dividing the net distance traveled during the cycle by the duration of the cycle. “Move” periods were defined as when the speed was above the 90th percentile, and “no move” periods were defined as when the speed was below the 10th percentile. Note that this measure of movement does not include running on the wheel but only moving through the T-maze. Recording sampling rate was 1252 Hz for neural data and 39.0625 Hz for position data. Prior to cycle-by-cycle analysis, a broad bandpass filter (1-25 Hz) was applied to these signals in order to improve extrema localization. During cycle segmentation the narrow bandpass filter cutoff frequencies were 4 and 10 Hz.

### Motor cortical beta recordings

Electrocorticography recordings from 23 patients with Parkinson’s disease were obtained during surgery for implantation of a deep brain stimulator and publicly released (de Hemptinne et al. 2015). Briefly, a strip of electrodes with 1 cm contacts was inserted over the primary motor cortex (M1) and re-referenced using a bipolar montage of adjacent contacts. The signals analyzed in this study were from a single channel in which one of the electrodes was over M1. Recordings were collected for 30 seconds before and during DBS. For more information on the data collection see de Hemptinne et al. (2015). Recording sampling rate was 1000 Hz. Prior to cycle-by-cycle analysis, signals were lowpass filtered at 200 Hz, and high frequency peaks (60 Hz and above) were removed (identical to Cole et al. (2017)). During cycle segmentation the narrow bandpass filter cutoff frequencies were 13 and 30 Hz.

### Visual cortical alpha recordings

The electroencephalography recording analyzed in Figure 10 was taken from a previously published study (Peterson and Voytek 2017). Briefly, the recording contained two sections. First, the subject was at rest with eyes closed for two minutes prior to beginning an experiment. Second, two minutes of data were analyzed during which the subject was performing a task in which they fixated and pressed a button when they perceived a faint visual stimulus. The trace analyzed was collected from position Oz, referenced to FCz. For more information on the data collection, see Peterson and Voytek (2017). Recording sampling rate was 500 Hz. Prior to cycle-by-cycle analysis, a broad bandpass filter (2-50 Hz) was applied to these signals in order to improve extrema localization. During cycle segmentation the narrow bandpass filter cutoff frequencies were 6 and 14 Hz.

## Results

### Cycle-by-cycle oscillatory feature distributions

In order to demonstrate the usefulness of our method, we applied our cycle-by-cycle analysis technique (Figure 1, see Methods) to recordings of theta oscillations in the pyramidal layer of CA1 and layer 3 of the entorhinal cortex (EC3) in a rat (Mizuseki et al. 2014).

Example traces from CA1 and EC3 are shown in Figure 2A-B, and the remaining panels summarize the oscillatory properties of these recordings. While the amplitude of the CA1 oscillation varies roughly between 1-2.5 mV, the theta oscillations in EC3 extend up to 4 mV (Figure 2C). However, the frequency of these two oscillators were very consistent with one another, as the distribution of periods mainly ranged between 100-150 ms, corresponding to 7-10 Hz (Figure 2D) in both CA1 and EC3. Theta oscillations in both regions exhibited reliable rise-decay asymmetry (rdsym) such that the rise period was systematically shorter than the decay period (Figure 2E). The peak around rdsym=0.4 represents that the rise period generally made up around 40% of the cycle, or in other words, that the decay period was about 50% longer than the rise period. The theta oscillation rdsym in CA1 was more variable than in EC3, shown by the broader distribution (Figure 2E, black). The CA1 and EC3 differed most in terms of their peak-trough symmetry (ptsym). Figure 2F shows that the CA1 theta oscillations tend to have shorter peaks than troughs (ptsym < 0.5) while EC3 theta oscillations tend to have shorter troughs and longer peaks (ptsym > 0.5). While ptsym is opposite between CA1 and EC3, this cannot be explained by a simple inversion in polarity, because that would require opposite rise-decay symmetries, which is not the case (Figure 2E).

It is interesting to consider the correlations between features in order to determine whether they contain redundant information. For example, cycle asymmetry may be unidimensional in theta oscillations, meaning that those cycles that are more rise-decay asymmetric are also the ones that are more peak-trough asymmetric. However, there is only a weak correlation between these two symmetries in the analyzed CA1 recording (Spearman correlation, r=-0.07). Additionally, there was a positive correlation between the rise-decay symmetry and the period (r=0.18), but there was no correlation between this symmetry of the theta oscillations and their amplitudes (r=0.0002). Note that p-values are not reported for this single-recording analysis because they should be interpreted with caution. The statistics assume that each cycle is independent, while in fact there are significant autocorrelations in cycle-by-cycle features (see below).

In order to restrict our oscillatory feature analysis to only the portions of the signal in an oscillatory regime, we applied an algorithm to determine which “cycles” that the original signal had been segmented into are truly part of an oscillation. We set requirements for the amplitude and periods of consecutive cycles to be similar, for flanks to be monotonic, and for a minimum number of cycles in each oscillatory period (Figure 3A-F, see Methods). These requirements can all be controlled by hyperparameters. For example, Figure 3G shows how increasing the amplitude and period consistency requirements decreases the fraction of the signal that is defined to be in an oscillatory regime (minimum monotonicity = 0.8, see Methods).

### Accuracy of detecting and characterizing oscillatory bursts

In order to assess the accuracy of our cycle-by-cycle oscillatory detection and characterization, we simulated a neural signal with a bursting oscillatory process and noise (Figure 4A-D, see Methods). This simulated signal had four times the power in the oscillation compared to the brown noise process (SNR=4). First, we tested if the algorithm can determine which periods of the signal were simulated in an oscillatory regime. Figure 4E-F shows the precision and recall of this algorithm with different amplitude and period consistency requirements. Intuitively, as the requirements become more strict (increase), a higher fraction of the cycles declared as oscillating are truly oscillating (high precision), but a lower fraction of the truly oscillating cycles are detected (low recall). The Fβ score is used to trade off between precision and recall, where β is the relative importance weight between precision and recall. For example, the F1 score (Figure 4G) weights precision and recall equally. For this signal, the F1 score is maximized when the amplitude consistency hyperparameter is set to 0.4 and the period consistency hyperparameter is set to 0.55. In this case, the algorithm’s precision is 75% and its recall is 82%. However, if we are more concerned that the cycles we analyze are truly oscillating than capturing all truly oscillating cycles, we can weight precision more than recall by setting β < 1.0. In this scenario, the Fβ score (β = 0.2) is maximized when the amplitude consistency hyperparameter is set to 0.6 and the period consistency hyperparameter is set to 0.75 (Figure 4H, precision = 97%, recall = 29%). That is, we are most accurate at determining oscillatory periods when we require adjacent cycles to be at least 60% similar (i.e., differ by no more than 40%) in amplitude and 75% similar in period. Note that the monotonicity requirement was fixed to 0.8 (see Methods).

Additionally, we assessed how well the properties of each cycle of the simulated oscillation matched the feature measurements after noise was introduced. Figures 4I-K show positive correlations between the simulated and measured values of amplitude (r=0.64), period (r=0.82), and rise-decay symmetry (r=0.67). The cycle amplitude tended to be overestimated because the measurement is positively biased by noise (Figure 4I). The cycles were also estimated to be more asymmetric than their true simulated structure (Figure 4K).

### Cycle-by-cycle vs. instantaneous measures of amplitude and frequency

We believe that measuring amplitude and frequency measures on a cycle-by-cycle basis could provide a better estimate of these properties compared to conventionally used instantaneous Hilbert-based measures, because the latter are not direct measurements of these properties but rather are based on filtering and nonlinear transformations. Therefore, we compared how these measures could differentiate conditions in a simulated experiment that elicit oscillations with differing amplitude and frequency (see Methods). This hypothetical experiment caused a 10 Hz oscillation after an event. There were three conditions: “baseline”, “higher amp”, and “longer bursts”. The oscillations in the “higher amp” condition were 20% greater in amplitude than in the other conditions. The “longer bursts” condition was 50% more likely to enter an oscillatory burst and 50% less likely to leave a burst relative to the other conditions. Instantaneous and cycle-by-cycle amplitudes were computed for the simulated trials (100 per condition) and averaged across each condition. The average instantaneous amplitude trace for “higher amp” was appropriately greater compared to the “baseline” condition (Figure 5A). However, the instantaneous amplitude for the “longer bursts” trials was even greater, even though these cycles had the same amplitude as the “baseline” trials. This undesirable trait is the result of averaging many non-stationary processes across many trials (Latimer et al. 2015; Jones 2016). In contrast, the cycle-by-cycle amplitude measure was appropriately increased specifically only for the “higher amp” condition (Figure 5B). Additionally, the cycle-by-cycle approach allows analysis of whether the signal is in an oscillatory burst, and so we can observe that the “longer bursts” condition was more commonly oscillating than the other two conditions (Figure 5C). This shows how cycle-by-cycle measures can capture features of oscillatory amplitude better than conventional Fourier-based methods.

Further, we compared how these two approaches to measuring oscillatory amplitude can statistically differentiate these conditions. To do this, the mean amplitude was computed between 500 and 1000 ms in each trial. The mean trial instantaneous amplitude only weakly differentiated the “baseline” and “higher amp” conditions (Figure 5D, U=4232, p=0.03) while the mean cycle amplitude more significantly differentiated these conditions (Figure 5E, U=123, p<10^-13^). This is due in large part to the ability of the cycle-by-cycle method to focus only on the portions of the signal in which the oscillation is present.

Similarly, we simulated conditions of this experiment that frequency measures should be able to differentiate. The “faster” condition simulates an 11 Hz (instead of 10 Hz) oscillation and the “nonsinusoidal” condition simulates an oscillation with relatively steep rises (rdsym = 0.2). The instantaneous frequency was averaged across trials for each condition (Figure 6A). Similarly, frequency was computed from the cycle-by-cycle period measures and averaged across trials (Figure 6B). Both Hilbert and cycle-by-cycle approaches were able to differentiate between the 10 Hz and 11 Hz oscillators (Figure 6C-D, instantaneous U=1800, p<10^-14^; cycle U=70, p<10^-15^). However, there was a spurious increase in instantaneous frequency in the “longer bursts” condition compared to the “baseline” condition (Figure 6E, U=3395, p<10^-4^). This is likely because instantaneous frequency is biased by aperiodic portions of the signal, depending on the distribution of power in the filtered frequency band, relative to periodic portions. In contrast, there was no difference in the cycle frequency measured in the “baseline” and “longer bursts” conditions (Figure 6F, U=1729, p=0.20). Both approaches correctly measured no frequency difference between the sinusoidal and nonsinusoidal oscillations (Figure 6G-H, instantaneous U=4939, p=0.44; cycle U=1017, p=0.20).

**Figure 6.**
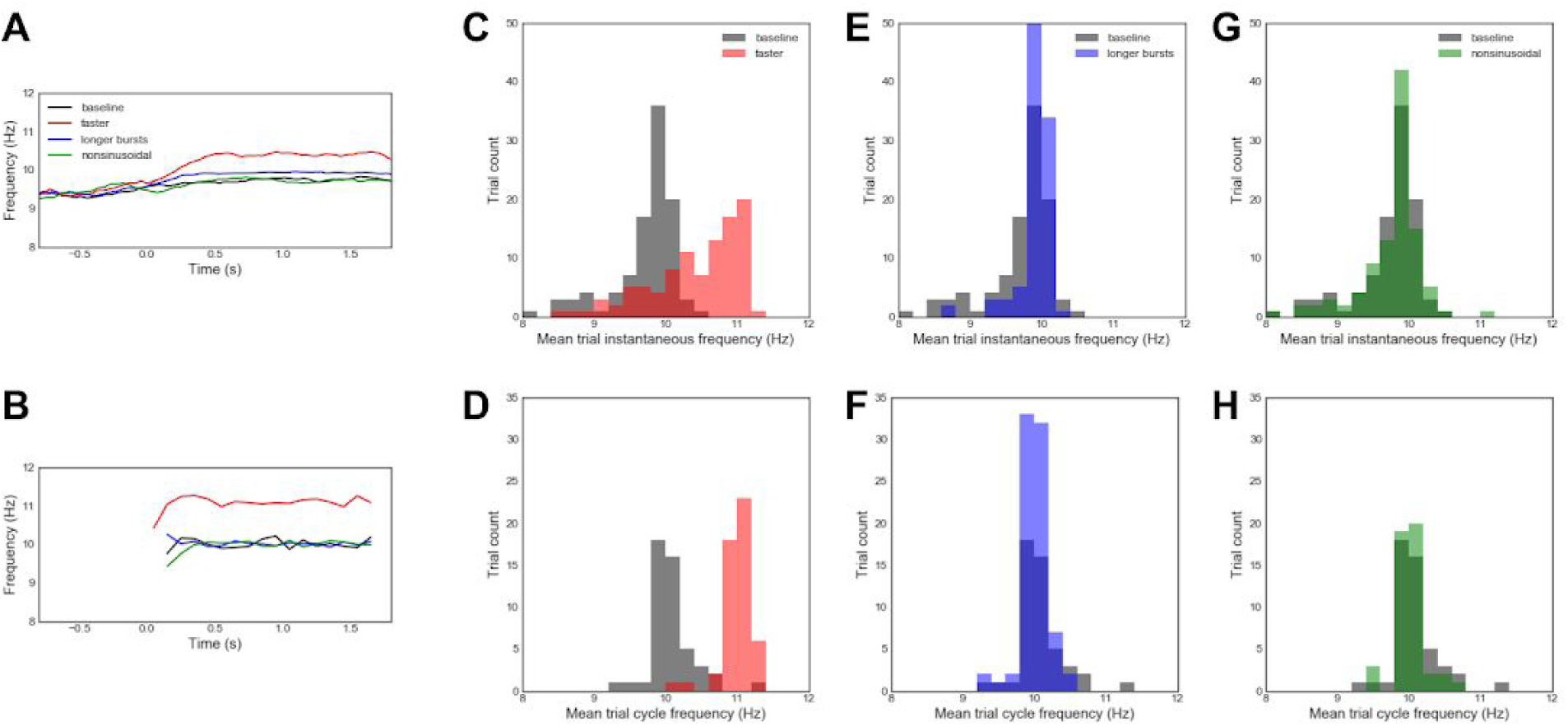
Comparison of cycle-by-cycle and instantaneous frequency analysis on simulated data. Four different event-related oscillatory changes were simulated for an event occuring at 0.0 seconds: (1,2) “baseline” and “longer bursts” conditions, similar to (Figure 4); (3) a “faster” condition wherein an 11 Hz oscillation was simulated instead of a 10 Hz oscillation, and; (4) a “nonsinusoidal” condition where oscillations had a constant rise-decay symmetry value of 0.2 (other conditions: 0.5). **(A)** Instantaneous Hilbert frequency profiles averaged across the four trial types: “baseline” (black), “faster” (red), “longer bursts” (blue), and “nonsinusoidal” (green). **(B)** Same as **(A)**, but the cycle-by-cycle frequency estimate was averaged across trials and binned at 100 ms intervals. This frequency estimate is not defined when no oscillating bursts are detected, hence the lack of an average before the event. **(C,D)** Distributions of the average **(C)** instantaneous frequency and **(D)** cycle frequency between 500 ms and 1000 ms for each trial in the “baseline” and “faster” conditions. **(E-H)** Same as panels C-D but comparing the “baseline” condition to the **(E,F)** “longer bursts” condition and the **(G,H)** “nonsinusoidal” oscillation condition. Note that conventional Fourier and Hilbert approaches capture event-related frequency increases **(A,C)** but also mischaracterize longer bursts as faster oscillations **(E)**. This mischaracterization is mitigated by the cycle-by-cycle approached introduced here, which successfully differentiates event-related changes in frequency from other conditions **(B,D,F,H)**.

### Autocorrelations of cycle features

An advantage of this cycle-by-cycle analysis framework is that it allows us to study the similarity of features between oscillatory cycles. For example, nearby cycles may have more similar symmetry properties compared to distant cycles. Through visual inspection, we observed that in nearby oscillatory bursts, the cycles in one may consistently have longer rise periods, whereas the cycles in the other burst may have relatively shorter rise periods. This would suggest that cycles within a burst have similar symmetry properties, which is shown by a significantly positive autocorrelation (Figure 7A). Additionally, nearby cycles are more likely to have similar amplitude and period, but these correlations are slightly confounded by the period and amplitude consistency requirements imposed in defining whether a cycle is to be considered for analysis. Therefore, we reran the analysis using an oscillatory burst detection algorithm based solely on an amplitude threshold (Feingold et al. 2015). Adjacent cycles remained correlated in their amplitudes (r=0.30), rise-decay symmetry (r=0.41), and peak-trough symmetry (r=0.23), but less so for period (r=0.06). Results were similar in another control analysis that selectively removed the amplitude (or period) consistency requirement and then assessed similarity between adjacent cycles (amplitude: r=0.48, period: r=-0.002).

**Figure 7.**
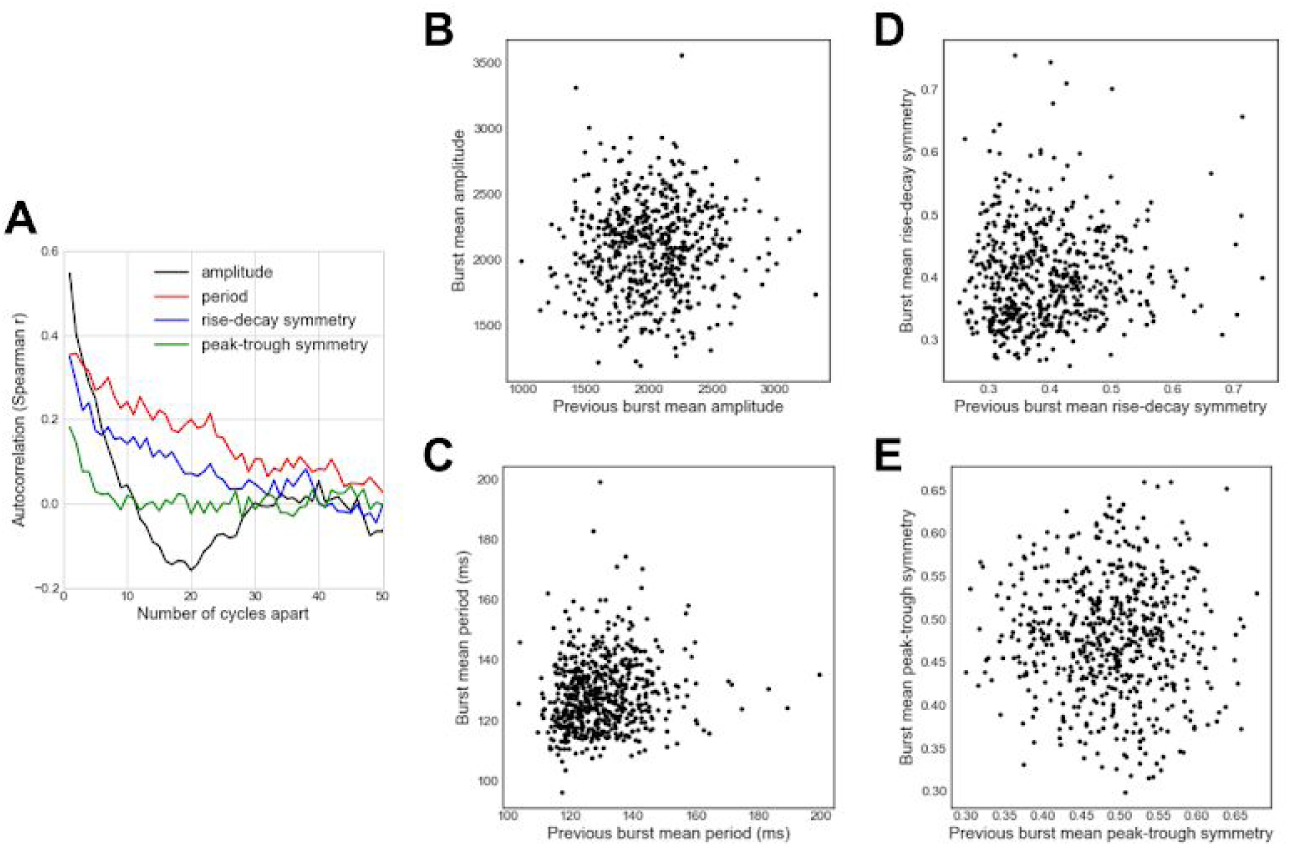
Autocorrelation of cycle features within, but not between, bursts of the hippocampal theta oscillation. **(A)** Autocorrelation of amplitude, period, and symmetry show that these features tend to be positively correlated for nearby cycles of the hippocampal theta oscillation recorded in an example rat. Note that the amplitude and peak-trough symmetry autocorrelations decay most rapidly. **(B-E)** The mean **(B)** amplitude, **(C)** period, **(D)** rise-decay symmetry, and **(E)** peak-trough symmetry of a burst are largely uncorrelated with the features of the previous burst. These results are not meant to conclusively demonstrate a physiological rule, but rather are used to highlight how novel information can be extracted using cycle-by-cycle approaches.

Because our analysis framework splits the recording into distinct oscillatory bursts, we also tested if consecutive bursts have correlated features. For this, we computed the burst feature (amplitude, period, or symmetry) as the mean of its cycles’ features. Whereas consecutive cycles had a strong positive correlation in their amplitudes, there was no correlation between the average amplitudes of consecutive bursts (Figure 7B, r=0.06, p=0.17). There was a modest correlation between the theta period (frequency) in adjacent bursts (Figure 7C, r=0.17, p<0.001), weak correlation between rise-decay symmetries (Figure 7D, r=0.09, p=0.03), and no correlation between peak-trough symmetries (Figure 7E, r=-0.02, p=0.65) of adjacent bursts. Together, this analysis suggests that the theta generative process during this recording is similar between cycles within a burst, but that there was a substantial change in the physiological parameters between oscillatory bursts.

### Cycle-by-cycle analysis to differentiate conditions

We demonstrate how the cycle-by-cycle analysis technique can be applied to three commonly analyzed oscillations: hippocampal theta, motor cortical beta, and visual cortical alpha. We compared a rat’s CA1 hippocampal theta oscillations between periods when it was moving and not moving (see Methods). Figures 8A-B show example recording segments in which the rat is moving and not moving, with its position plotted in Figure 8C. Figure 8D-G compare the theta oscillatory features between cycles in which its measured speed was below the 10th percentile (not moving, 1430 cycles), and cycles in which the measured speed was above the 90th percentile (moving, 1430 cycles). There was a slight increase in theta amplitude with running (Figure 8D) and a notable decrease in period (Figure 8E), as previously reported (McFarland et al. 1975; Sławińska and Kasicki 1998). Additionally, previous studies have qualitatively reported that theta oscillations are more rise-decay asymmetric (shorter rise) when running (Buzsáki et al. 1985; Belluscio et al. 2012; Hentschke et al. 2007), and we quantify that effect here (Figure 8F), as well as identify an increase in peak-trough asymmetry (Figure 8G, shorter peak). Additionally, scatter plots in Figures 8H-K show these effects when considering each individual cycle. Most notably in these plots, the time periods with the highest speed (*log*(speed) > 1.5 a.u.) tend to have different feature distributions compared to the cycles at which the rat was running at lower speeds.

**Figure 8.**
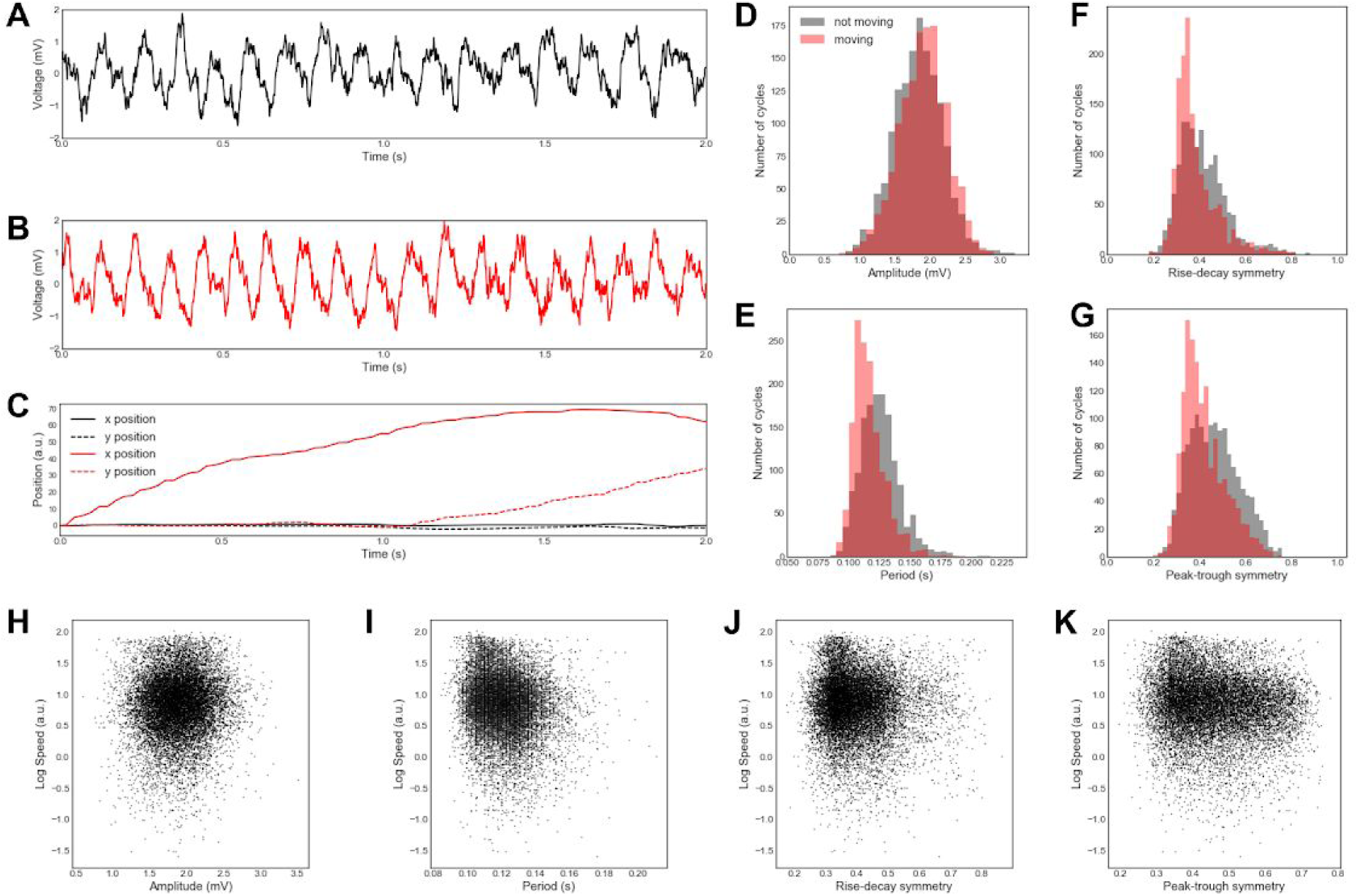
Difference between hippocampal theta oscillations between running and not running in an example rat. **(A-B)** Raw field potential in CA1 pyramidal layer during a period of **(A)** not running and **(B)** running during the same recording. **(C)** Position of the rat at rest (black) and while running (red). **(D-G)** Comparison of **(D)** amplitude, **(E)** period, **(F)** rise-decay symmetry, and **(G)** peak-trough symmetry of theta oscillations in the same example recording during running (red) and not running (black). Our approach shows in this recording that running is **(D)** not associated with a change in theta amplitude, but is associated with, **(E)** a shorter period (faster frequency) and, **(F,G)** greater asymmetry, as previously reported. **(H-K)** Correlations between the average speed of a rat in a cycle and the **(H)** amplitude (r=0.05, p<10^-8^), **(I)** period (r=-0.15, p<10^-77^), **(J)** rise-decay symmetry (r=0.06, p<10^-11^), and **(K)** peak-trough symmetry (r=-0.11, p<10^-41^). Note that, in this example, the symmetry measures are physiological correlates of behavior **(K)**.

Next, we re-analyzed motor cortical recordings from a previous study in which it was shown that “sharpness asymmetry” of motor cortical beta oscillations was decreased with deep brain stimulation (DBS) treatment of Parkinson’s disease (Cole et al. 2017). The current “peak-trough asymmetry” measure was designed to measure the same intuitive sense of “sharpness,” but it differs in that it is computed as a temporal ratio that is independent of amplitude (Figure 1G, see Methods), whereas the previously used “sharpness asymmetry” was confounded with amplitude. Figure 9A-B show recordings from an example subject before and during DBS. In this subject, DBS decreased the amplitude of beta oscillations (Figure 9C), but did not affect their period (Figure 9D) or rise-decay symmetry (Figure 9E). Across the patient population (N=23), there was no consistent effect of DBS on amplitude (Wilcoxon Signed Rank test, W=104, p = 0.30), period (W=111, p=0.41), or rise-decay asymmetry (W=104, p=0.30). However, DBS did elongate the relative peak time in the example subjects(Figure 9F), and consistently caused the beta oscillations to become more peak-trough symmetric (Figure 9G, W=65, p=0.026), consistent with the previously published sharpness ratio results (Cole et al. 2017). Note that peak-trough asymmetry here is measured as the difference from a symmetric oscillation, such that 0 represents equal duration peaks and troughs, and 0.1 represents an oscillation in which the average cycle was 60% peak or 60% trough. This was done because the polarity was not consistent across recordings.

**Figure 9.**
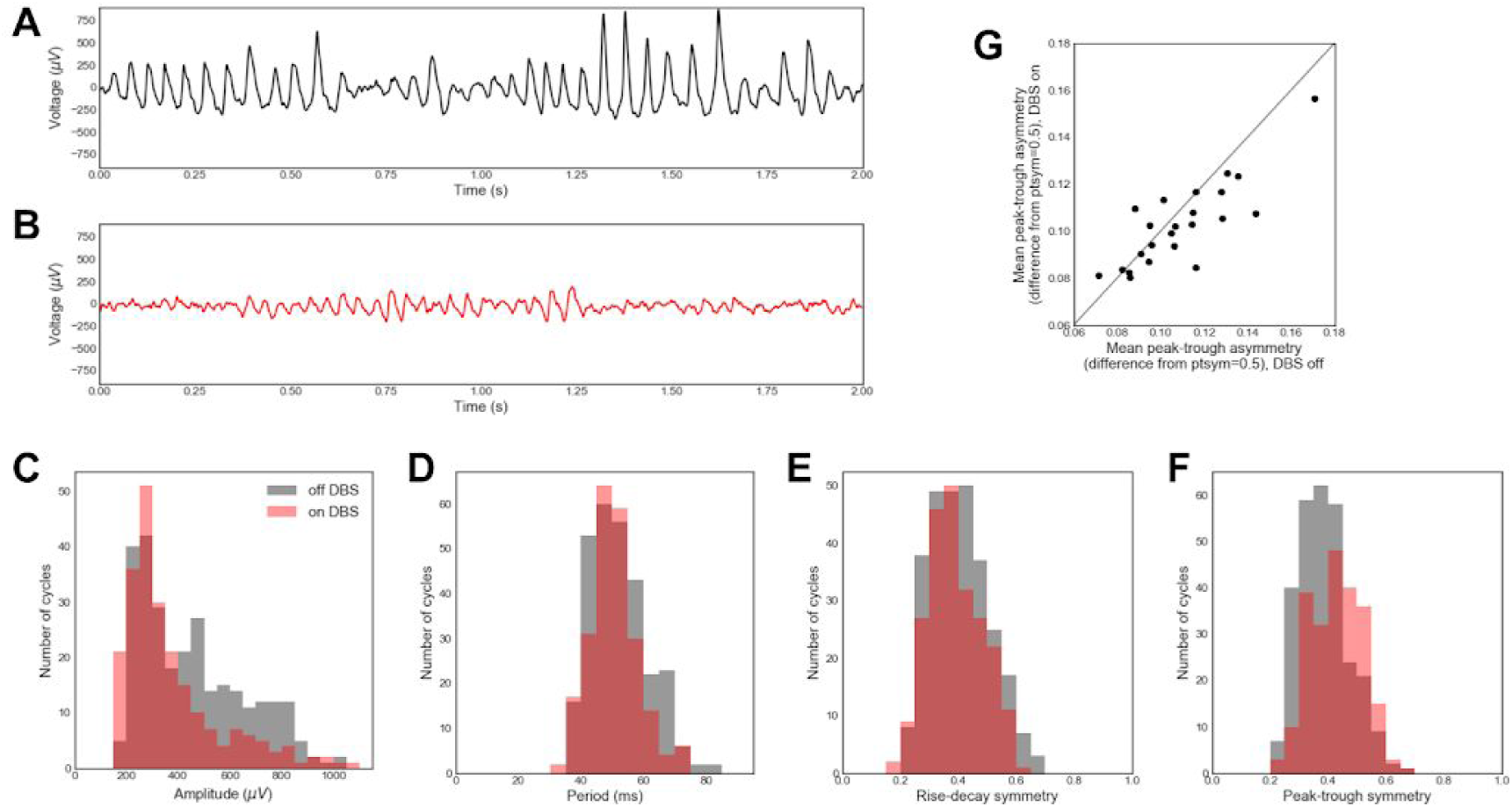
Changes in motor cortical beta oscillation shape with deep brain stimulation (DBS) treatment of Parkinson’s disease. **(A,B)** Motor cortical electrocorticography recordings from one subject (A) before and (B) during DBS. **(C-F)** Comparison of **(C)** amplitude, **(D)** period, **(E)** rise-decay symmetry, and **(F)** peak-trough symmetry of beta oscillations in the same subject before (black) and during (red) DBS. **(G)** Comparison of peak-trough asymmetry of beta oscillations before and during DBS. This value is computed as the absolute difference between the average peak-trough symmetry and 0.5 (equal peak and trough length). Each dot represents one subject. The diagonal line represents the same peak-trough asymmetry before and during DBS. These results extend previous work, showing that peak-trough symmetry, specifically during bursts, is reduced after DBS treatment in most patients.

Lastly, raw EEG recordings are shown for a subject while resting with eyes closed (Figure 10A) and while performing a visual target detection task (Figure 10B). We replicate the well-known observation that alpha oscillations are larger in the visual cortex while eyes are closed (Figure 10C). However, we do not observe a substantial difference in period, rise-decay symmetry, or peak-trough symmetry (Figure 10D-F). We do note that these distributions are more broadly distributed while the subjects’ eyes are open. This may reflect that there is more variance in the physiological parameters that govern the alpha oscillatory generator. However, this could also be attributed to a greater fraction of the alpha oscillations detected in the task condition actually being noise.

**Figure 10.**
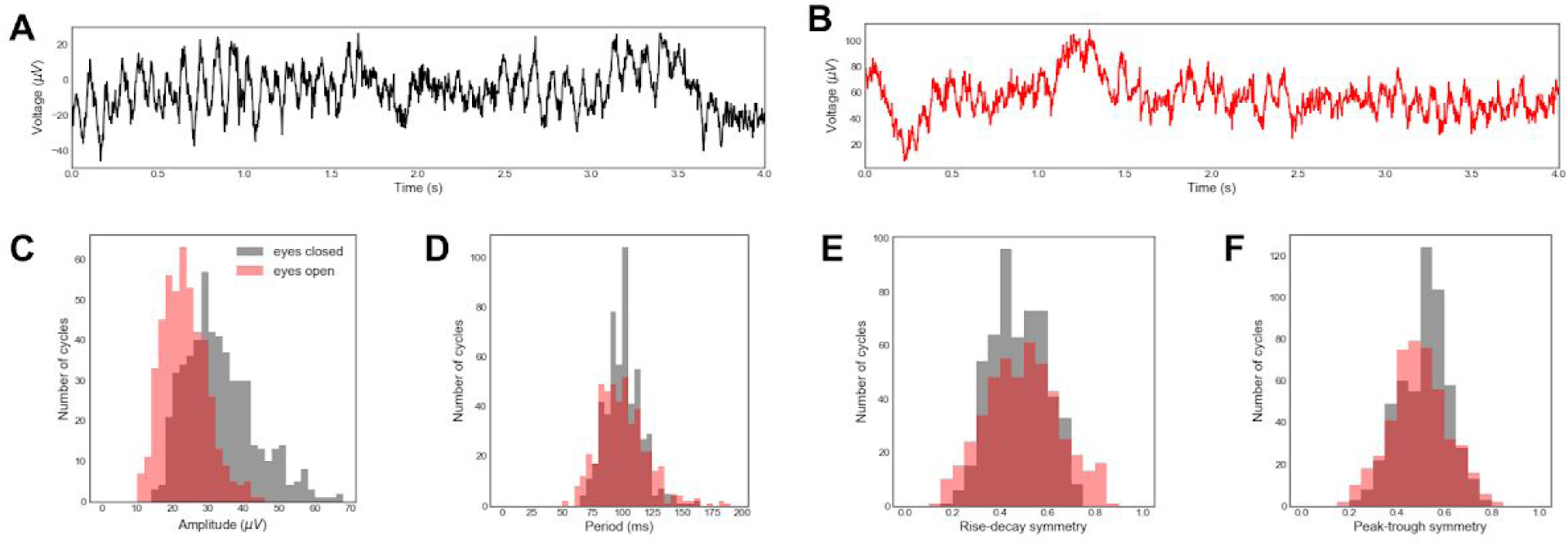
Differences in the visual alpha rhythm between periods of eyes-closed resting and eyes-open task in an example subject. **(A,B)** Example raw EEG traces while **(A)** the subject’s eyes were closed and **(B)** the subject was performing a task with eyes open. **(C-F)** Comparison of **(C)** amplitude, **(D)** period, **(E)** rise-decay symmetry, and **(F)** peak-trough symmetry of alpha oscillations between recordings with eyes closed (black) and open (red). Note that in this case, the well-known effect of eyes open/closed on alpha amplitude is recapitulated **(C)**, though eyes open/closed has no effect on period (frequency) **(D)**, rise-deay symmetry **(E)**, or peak-trough symmetry **(F)**. This demonstrates how cycle-by-cycle waveform analysis is complementary to Fourier-based approaches.

## Discussion

We have described a new technique for analyzing neural signals that has advantages over conventionally used signal processing techniques. Specifically, this cycle-by-cycle analysis approach quantifies waveform symmetry and restricts analysis to only periods in which an oscillation is present. This technique offers further analytic possibilities, beyond what is demonstrated in this paper. For example, rather than comparing cycles from separate recordings (e.g., DBS on vs. DBS off), cycles can be assigned to a trial and aggregated in order to analyze the effects of task conditions or correlates to responses, such as reaction time. Though we only covered four cycle features in this paper (amplitude, period, rise-decay symmetry, and peak-trough symmetry), additional features can be designed and easily added to this workflow, such as monotonicity of the flanks or gamma power. Additionally, the oscillatory detection algorithm allows for quantifying bursting features such as burst duration or burst rate that may correlate meaningfully to experimental parameters.

### Caveats of cycle-by-cycle analysis

Like Fourier-based analysis, there are also caveats of this cycle-by-cycle technique that need to be considered to minimize confounds. Because oscillatory features tend to be autocorrelated (Figure 7A), it will usually be invalid to treat each cycle as independent in statistical tests. This is a similar caveat to trial-wise analyses, in which consecutive trials are often not independent. To bypass this issue and assess significance within a recording, the recording can be split into multiple non-overlapping segments, and a statistical test can be performed on a metric of each segment (e.g., mean difference in rdsym between conditions A and B). It is also important to keep in mind that these cycle features are not independent of one another. This can be assessed, for example, by quantifying the correlation between cycle amplitude and rise-decay symmetry to potentially uncover that higher magnitude oscillations tend to be more asymmetric, and so these features may capture redundant information. In order to tease apart some interdependencies, multiple features could be incorporated into a model to predict a condition or behavior of interest (e.g., general linear model or logistic regression), and the unique contribution of each feature can be assessed. For example, if oscillatory amplitude is greater in condition A vs. B, and amplitude is correlated with symmetry, then we will also observe that symmetry differs between conditions. However, a multidimensional model could detect whether symmetry contains any additional information beyond that provided by amplitude.

One key advantage of this approach is that it is designed in a way that does not favor any specific oscillatory waveform (e.g., sinusoid). However, it is important to note that this approach is not entirely free from reliance on Fourier-based methods. The main instance of this is that a narrow bandpass filter is used in the process to localize oscillatory peaks and troughs (Figure 1C). Importantly, statistics are not ultimately computed on this narrowband signal, as it is only a tool that is useful for estimating peaks and trough times. Another instance of using Fourier-based techniques in this approach is the optional broad bandpass filter (Figure 1B). This is done in order to remove the slow transients or high frequencies that may make extrema identification more difficult. However, the filter is potentially removing some of the oscillatory signal that is outside of the broad passband. Therefore, the cutoff frequencies of the broad bandpass filter are chosen to compromise a tradeoff between the fidelity of waveform shape and the ability to localize peaks and troughs (Figure 1A-B).

Because extrema localization is nontrivial, caution is necessary when performing analysis that considers the precise times of peaks and troughs. For example, an aperiodic process could delay the algorithm’s trough localization, and so if a neuron fires most at the trough, it appears that it fires at an earlier phase when the decay period is artificially elongated. However, Belluscio et al. (2012) reported that a rat’s position could be better decoded using extrema interpolation (Figure 1F, black) compared to the conventional Hilbert transform-based method (Figure 1F, red). If it is difficult to filter the signal in order to achieve reasonable extrema localization and symmetry fidelity, then the oscillation may not be suitable for cycle-by-cycle analysis. For example, it is likely not reasonable to analyze the beta frequency band in the visual cortex because the presence of this rhythm is usually not evident in the time series, whereas alpha is prominent.

Hyperparameter selection is another notable challenge. Conventional analysis forces semi-arbitrary choices during filter design, which can have significant impacts on results. In the cycle-by-cycle approach, the user must also define the specific narrowband frequency range of interest and also whether to divide cycles peak-to-peak or trough-to-trough. We recommend that the user runs the analysis with multiple hyperparameter choices in order to test for robustness of their results. Setting the thresholds for defining oscillatory bursts requires considerable parameter selection. For analysis of motor cortical beta and visual cortical alpha rhythms in this paper, thresholds were tuned while visualizing the algorithm’s output until the oscillatory burst detection seemed most accurate.

### Cycle-by-cycle approach to defining bursts of oscillations

A key feature of this oscillatory burst detection method is that it does not need to set an amplitude threshold to define bursting periods, as is the case for previously published algorithms for burst detection (Hughes et al. 2012; Feingold et al. 2015; Watrous et al. 2017). This makes the current burst detection algorithm especially suitable for detecting oscillators that may occur at both small and large amplitudes, or may vary greatly in stationarity between recordings. In contrast using an amplitude threshold based on scaling the median oscillatory power (Feingold et al. 2015), inherently defines an upper limit on the fraction of the signal that can be oscillatory, which may not be suitable in a hippocampal recording that oscillates at a theta frequency sometimes over 50% of the time.

It should be noted that the optimal hyperparameters for the current algorithm to maximize accuracy (Figure 4) are heavily dependent on the parameters chosen for the simulated data. For example, while the F1 score is optimized for the current signal (SNR=4) when the amplitude consistency requirement was set to 0.4 and the period consistency requirement was set to 0.55, if the signal SNR is decreased to 1, an amplitude consistency requirement of 0.3 and period consistency requirement of 0.5 optimizes F1 score. In this case, less strict requirements balanced precision and recall, likely because true oscillatory regimes are inherently more difficult to detect in the higher noise scenario. Additionally, the correlations between simulated and measured cycle features were decreased when SNR was lowered (amplitude: r = 0.34, period: r = 0.70, rdsym: r = 0.45). This implies that more cycles are necessary in order to obtain an accurate estimate of cycle features when the SNR is decreased.

Very weak correlations between cycle features of adjacent bursts (Figure 7B-E) was a particularly surprising result. One could have expected there to be a larger correlation between features of adjacent bursts compared to features of adjacent cycles because individual cycles may be noisy estimates, and averages may lead to more accurate estimates. However, these correlations between adjacent bursts were very slight, if at all present, compared to the correlations between features of adjacent cycles. This suggests that the properties of this rat’s hippocampal theta oscillation are consistent within an oscillatory burst but vary considerably from burst to burst. However, results were qualitatively different when this analyzing a recording from another rat, and so this phenomena should be investigated in future, more controlled, studies.

Unlike amplitude and symmetry, we did not observe a positive correlation between the periods of adjacent cycles after controlling for the period consistency requirement of the burst detection algorithm. This is likely due to the effect of noisy extrema estimation on the period estimate. For example, if noise in the signal causes a cycle’s latter trough to appear delayed, the current cycle would be measured as artificially longer, and the subsequent cycle would appear artificially shorter. This would produce a negative correlation between the periods of adjacent cycles. However, we observe no correlation between consecutive periods, so it is possible that this effect is masking a positive correlation.

### Instantaneous and cycle-by-cycle measure comparison

In addition to its ability to quantify waveform symmetry, we believe that the cycle-by-cycle framework’s measures of amplitude and period also offer an advantage over current instantaneous estimates of amplitude and frequency. An important step in computing instantaneous amplitude is convolution with a kernel of the frequency of interest. This means that the amplitude measure at any given point in time is actually computed using data from several cycles around that point (depending on the filter length). Because convolution is a linear operation, it is not specifically sensitive to oscillatory amplitude, but it will be strongly biased by non-oscillatory sharp transients. Instantaneous frequency, derived from instantaneous phase, is similarly based on this convolution, and fluctuates within a cycle due to the cycle’s temporal dynamics. However, when applied to a relatively stationary nonsinusoidal oscillation (e.g., hippocampal theta) this will cause fluctuating within-cycle frequency estimates, which do not actually reflect a change in the theta frequency, but rather reflects its sawtooth-like waveform.

In contrast to these widely used instantaneous measures, the time-resolved estimates of amplitude and period (frequency) using cycle-by-cycle estimates are more direct and intuitive measurements of the oscillator. Specifically, the amplitude measures the mean rise and decay voltages, and the period is computed as the time between consecutive peaks of a cycle in a putative oscillatory burst. This method does not over-promise temporal resolution that it cannot reliably account for, and it is robust to issues that plague instantaneous measures like sharp transients and nonsinusoidal waveforms. Additionally, we showed that the cycle-by-cycle measures of amplitude and frequency are more robust and better at differentiating these properties in simulated oscillations (Figures 5-6). Specifically, both of these instantaneous measures are biased by the proportion of the signal in which the oscillator is present, so this could underlie some past reports of changes in instantaneous amplitude and frequency.

There is some precedence for analyzing individual cycles of brain rhythms. Several researchers studying the hippocampal theta rhythm have quantified its rise-decay asymmetry by measuring the rise and decay times of each individual cycle (Belluscio et al. 2012; Dvorak and Fenton 2014). Additionally, hippocampal theta researchers have designed an alternative instantaneous phase estimate that involves identifying extrema in each cycle (Siapas et al. 2005; Belluscio et al. 2012). Much earlier, Adrian and Matthews (1934) studied the evolution of a gamma oscillation in response to injury to the cortex of a cat. Initially, they observed rhythmic transient discharges, which gradually became more frequent and broad, producing a more sinusoidal-like rhythm. They interpreted the initial transients as bursts of activity by a few local neurons, and that this activity spread out as the transient discharges merged into a quasi-sinusoid. Therefore, each cycle could be considered as a “packet” of neural activity that can be characterized distinctly from the previous and subsequent cycles using a cycle-by-cycle analysis framework. This view of each cycle as an informative physiological unit differs substantially from modern work on oscillations.

### Conclusions

In summary, we have demonstrated a novel approach to analyzing neural oscillations using a cycle-by-cycle framework. This technique has advantages over conventional Fourier-based techniques including its characterization of oscillatory waveform symmetry and inherent detection of oscillatory regimes. We demonstrate its applicability to hippocampal theta, motor cortical beta, and visual cortical alpha rhythms. Applications are not limited to the oscillations discussed here but could also include other prominent rhythms such as sensorimotor mu, visual cortical gamma, thalamocortical spindles, cortical slow oscillation, and respiratory rhythms. While this open-source analysis framework is unique in its focus on oscillatory symmetry, it is also complementary to conventional analysis of oscillatory amplitude and period, and, as such, should be a standard part of the neural oscillation analysis toolbox.

## Acknowledgements

We thank Richard Gao, Tammy Tran, and Tom Donoghue for invaluable discussion and comments on the manuscript. S.R.C. is supported by the National Science Foundation Graduate Research Fellowship Program and the University of California, San Diego Chancellor’s Research Excellence Scholarship. B.V is supported by a Sloan Research Fellowship, the Whitehall Foundation (2017-12-73), and the National Science Foundation (1736028).

## Footnotes

1 https://pypi.python.org/pypi/neurodsp/0.3

2 https://github.com/voytekresearch/neurodsp

